# Single-cell multiomics reveals the distinct properties of neonatal and adult recent thymic emigrants

**DOI:** 10.1101/2025.02.26.640106

**Authors:** Cybelle Tabilas, Vanessa Venturi, Connor Kean, Anastasia Minervina, Paul G. Thomas, Andrew Grimson, Jennifer K. Grenier, Miles P. Davenport, Norah L. Smith, Brian D. Rudd

## Abstract

Following thymic egress, CD8+ T cells must undergo a post-thymic maturation process to transition from a recent thymic emigrant (RTE) to a mature naïve T cell. Since the neonatal T cell pool is comprised of significantly more RTEs, the prevailing notion is that neonatal CD8+ T cells behave differently than their adult counterparts simply because they have undergone less post-thymic maturation. To test this theory, we leveraged a fate mapping mouse model and paired single cell transcriptome and TCR sequencing to compare neonatal and adult CD8+ RTEs that have undergone the same amount of post-thymic maturation. Interestingly, we found that neonatal and adult CD8+ RTEs exhibit distinct phenotypes, gene expression profiles, TCR usage, and functions. These data suggest that neonatal CD8+ T cells are not simply immature adult CD8+ T cells and that age-related changes in CD8+ T cell functions in early life cannot be attributed solely to differences in the amount of post-thymic maturation.

## Introduction

Previous work has shown that the phenotype and function of CD8+ T cells continue to evolve after their egress from the thymus ^1-5^. The subset of murine CD8+ T cells that are less than 3 weeks old, denoted recent thymic emigrants (RTEs), are phenotypically and functionally distinct from their more mature counterparts ^6,7^. While RTEs make up a small percentage (∼10-20%) of naïve cells in adult mice, they account for nearly 100% of T cells in neonatal mice ^8^. Interestingly, the functional differences between RTEs and mature T cells in adult mice are reminiscent of the differences that have been previously described in naïve CD8+ T cells from neonatal and adult mice. For example, both adult CD8+ RTEs and neonatal CD8+ T cells preferentially become short-lived effectors during infection ^4,9-11^. Moreover, adult CD8+ RTEs and neonatal CD8+ T cells exhibit more innate-like functions, as evidenced by their innate-like receptor expression (complement receptors and NK cell receptors) and ability to deploy non-specific defense mechanisms more typically associated with innate cells ^12-19^. As a result, there is an assumption that neonatal CD8+ T cells are simply immature adult CD8+ T cells or adult CD8+ RTEs.

To test the theory that developmental changes in the CD8+ T cell response correspond to differences in post-thymic maturation, it is important to directly compare RTEs from neonatal and adult mice. If neonatal CD8+ T cells behave differently than adult CD8+ T cells because they are comprised of more RTEs, then age-related differences in CD8+ T cells should disappear when we directly compare neonatal and adult RTEs. Unfortunately, comparing neonatal and adult CD8+ RTEs is challenging because there is currently no marker to distinguish RTEs from mature CD8+ T cells, and the experimental strategies that have been developed to label RTEs in mice have well-known limitations. For example, injection of fluorescein isothiocyanate (FITC) into the thymus can mark a small number thymocytes that later become FITC+ RTEs in the periphery, but the introduction of surgical stress may alter the behavior of RTEs ^20,21^. Others have used BrdU incorporation to label the rapidly dividing thymocytes before they are exported into the periphery as RTEs, but BrdU can also be incorporated into mature T cells that are undergoing cell division ^22,23^. Perhaps the most useful method for identifying RTEs is to use Rag2p-GFP transgenic (Tg) mice, whereby GFP expression remains detectable in peripheral T cells for a finite period of time (∼3 wks), thereby serving as a marker for RTEs ^4,5,24^. However, the major disadvantage of this approach is that the GFP signal is lost in cells that undergo extensive homeostatic proliferation, making them unsuitable for identifying RTEs in the more lymphopenic neonatal mice ^25,26^.

In this study, we used a new method to identify RTEs in both neonatal and adult mice. Our approach involved the use of fate-mapping mice to permanently label, or ‘timestamp,’ a wave of thymic CD8+ T cells at the time of tamoxifen exposure ^14,27,28^. The advantage of this approach is that we can identify neonatal and adult RTEs in the absence of surgical stress, and the label is not diluted out by proliferation. Along with timestamp mice, we employed the 10X Genomics Chromium Single Cell Immune Profiling platform (paired TCR/RNA-seq) to rigorously compare how the composition of RTEs differs in neonatal and adult mice, together with flow cytometry to assess their phenotype and functions. These experiments demonstrated that all CD8+ RTEs are not created equally, and the altered numbers of RTEs in neonatal and adult animals is not a major contributor to the age-related differences in CD8+ T cell behavior.

## Results

### Phenotypes of neonatal and adult CD8+ RTEs

The goal of this study was to determine whether CD8+ RTEs in neonatal and adult mice are phenotypically and functionally distinct. To accomplish this goal, we used a fate-mapping mouse model that allows us to permanently label RTEs produced at different ages. This model is based on the TCRδ-CreER strain ^29,30^. The TCRδ gene is expressed by all T cells in the thymus at the double negative (DN) and double positive (DP) stages of development but is excised in αβ T cells during their transition to the single positive (SP) stage of thymopoiesis ^31,32^. Thus, by crossing the TCRδ-creERT2 mice with a reporter strain that has a loxP-flanked ‘stop’ cassette upstream of a fluorescent reporter in the Rosa26 locus (R26R^Zsgreen^ or R26R^TdTomato^), we can excise the ‘stop’ cassette and permanently label, or ‘timestamp’, a wave of CD8+ T cells made in the thymus only during the time of tamoxifen exposure. We administered tamoxifen to one group of mice at 1d of age to mark a wave of neonatal CD8+ T cells and another group of mice at 28d of age to label a wave of adult CD8+ T cells (Fig. 1A). Two weeks after labeling, we collected timestamped CD8+ RTEs from the spleens of both groups of mice and compared their phenotype using flow cytometry.

**Figure 1.**
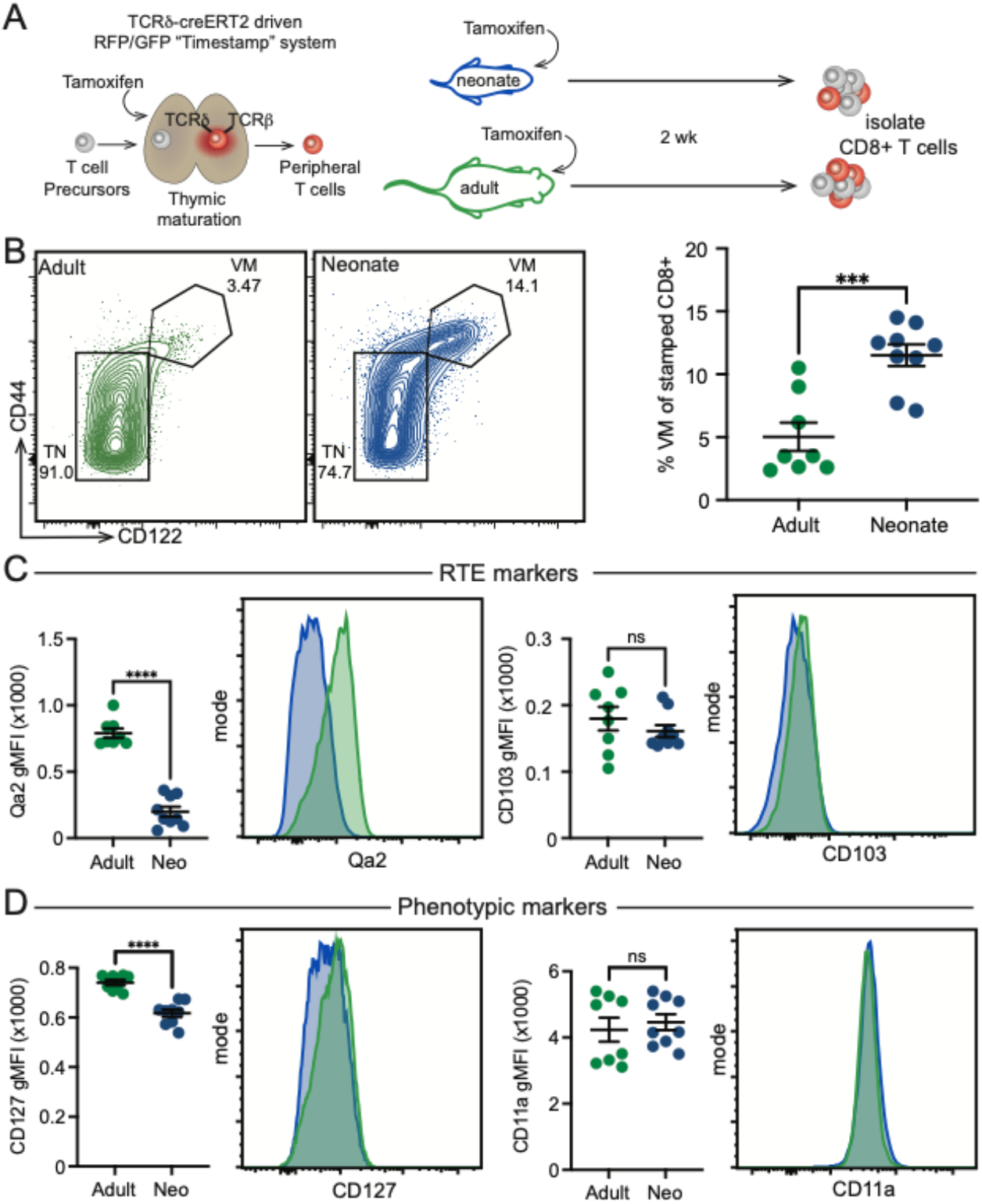
Neonatal and adult RTEs are phenotypically distinct. (A) Schematic of timestamping system marking T cells made at the time of tamoxifen administration. RTEs were collected at 2 weeks post marking. (B) Representative contour plots displaying virtual memory (VM, CD44^hi^ CD 122^hi^) and true naïve (TN, CD44^lo^ CD122^lo^) CD8+ populations (left) and statistical analysis of VM population within the marked T cell population (right). (C) Statistical analysis and representative histograms of RTE markers Qa2 (left) and CD103 (right). (D) Statistical analysis and representative histograms of phenotypic markers CD127 (left) and CD11a (right). N=9 from 2 independent experiments. For statistical analysis, unpaired t-tests were performed, ns, not significant; ***, P<0.001; ^****^ P<0.0001.

We previously reported that a large proportion of CD8+ T cells in neonatal mice exhibit a virtual memory (VM) phenotype (CD44^hi^CD122^hi^), whereas only a small fraction CD8+ T cells in adult mice show this phenotype ^10,33^. Thus, we were interested in determining whether this phenotypic difference would still be evident in neonatal and adult CD8+ RTEs that had undergone the same amount of post-thymic maturation. Interestingly, the neonatal CD8+ RTEs contained significantly more VM cells than their adult counterparts (Fig. 1B). The neonatal and adult RTEs expressed different amounts of RTE maturation marker Qa2 but similar levels of CD103 (Fig. 1C). Neonatal RTEs also had lower expression of the IL7 receptor CD127 but similar levels of CD11a (Fig. 1D). Thus, many surface markers that have traditionally been used to identify immature naïve T cells do not apply to CD8+ T cells produced in early life.

We considered the possibility that some of the phenotypic differences between neonatal and adult RTEs were driven by changes in the peripheral environment. In particular, neonatal CD8 RTEs are exported into a more lymphopenic environment, which may promote homeostatic proliferation and accumulation of virtual memory cells ^25,26^. To control for age-related differences in the peripheral environment, we used a dual-reporter timestamp system in which a thymic lobe from a 1-day-old TdTomato+ timestamp animal is transplanted under the kidney capsule of a 6-week-old Zsgreen+ timestamp animal (Fig. S1A). In this way, we could compare the phenotypes of neonatal and adult CD8+ RTEs in the same peripheral environment. Consistent with our findings in different-aged mice, we found that neonatal CD8+ RTEs still acquire a unique phenotype even when matured in a lymphoreplete adult environment (Fig. S1B-C). Furthermore, we found that neonatal RTEs show higher expression of the transcription factor Eomes (Fig. S1D) and are more proliferative, as indicated by an increased population that is Ki67+ (Fig. S1E). Collectively, these results suggest that phenotypic differences between bulk neonatal and adult CD8+ T cells cannot be solely attributed to differences in the amounts of post-thymic maturation.

### Gene expression profiles of neonatal and adult CD8+ RTEs

To understand how the composition and function of the RTE pool changes at different stages of life, we examined the single cell gene expression profiles of RTEs produced within neonatal and adult mice. We employed paired single cell RNA-seq (scRNA-seq) and single cell TCR-seq (scTCR-seq) to obtain both gene expression and TCR from individual CD8+ T cells ‘stamped’ in neonatal and adult mice. The stamped CD8+ T cells were sorted from the spleens of neonatal and adult mice at two weeks post stamp, allowing us to examine gene expression profiles in the same two groups of RTEs that were analyzed by flow cytometry.

To characterize the heterogeneity of the RTE pool, we performed dimensionality reduction and clustering of the 59,281 single cell transcriptome profiles using Seurat ^34^ (Fig. 2A). Having identified 11 total clusters, we then assigned functional annotations to each cluster considering both the cluster-specific upregulation of individual genes (Fig. 2B) and gene set module scores, a method that reports the expression of multiple genes within a pathway of interest at single cell resolution ^35^ (Fig. 2C). Broadly, the 11 clusters partitioned into three functionally distinct superclusters: 1) effector-like, 2) true naïve-like, and 3) virtual memory-like. Clusters 1 and 8 made up the effector-like supercluster, expressing genes typically upregulated post activation (Fig. 2B - *Nme1, Srm, Shmt1*) and having transcriptomes with high gene set module scores for “preparation for cell division” (Fig. 2C). Clusters 0, 3, 5, 6, and 9 made up the true-naïve-like supercluster, expressing genes associated with a naïve state (Fig. 2B – *Tsc22d3, Ccr9*) ^36,37^ and possessing transcriptomes with the highest “true naïve” gene set module scores. (Fig. 2C). Conversely, clusters 2, 4, 7, and 10 made up the virtual memory-like supercluster, expressing genes associated with a virtual memory phenotype (Fig. 2B – *Smad3, Tbx21, Ly6c2, Cxcr3*) ^14^ and having transcriptomes with the highest “virtual memory” gene set module scores (Fig. 2C).

**Figure 2.**
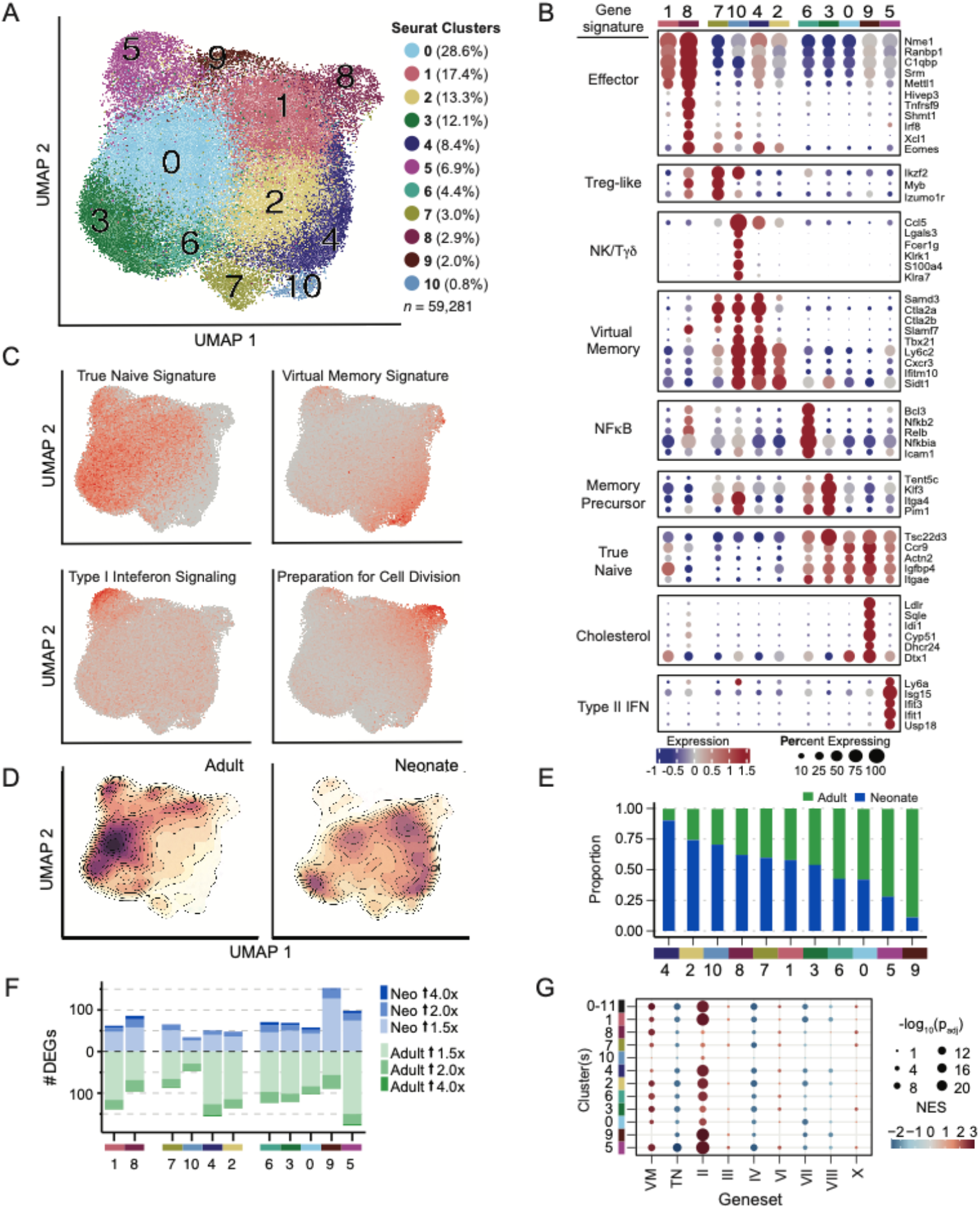
RTE pools possess age-related differences in composition and gene expression. (A) UMAP of neonate (32,185) and adult (27,096) RTE transcriptomes. Cluster identities are indicated by color, and cluster size is conveyed as a percentage of the total number of RTEs sequenced (59,281). N = 4 mice for both neonatal and adult groups. (B) Bubble plot depicting the expression of the top 5 marker genes. Genes are grouped together along the y- axis based on their average expression values across each gene expression cluster. Gene signature labels annotate the gene groupings. Dot shading indicates the mean normalized expression (row-wise z-score), and dot sizes represent the percentage of cells expressing each gene. (C) UMAP visualization of defined gene set module scores. Single cell gene set module scores are conveyed by point color. (D) Density estimation of single cells within the UMAP projection, split by age. Filled contour lines indicate regions of the UMAP with high and low densities of cells. (E) Proportion of adult and neonatal derived RTEs that compose each cluster normalized to the total number of adult and neonatal RTEs profiled. Clusters ordered from highest to lowest proportion of neonatal cells. (F) Number of age-related differentially expressed genes (DEGs) per cluster. Fill color indicates the fold-change magnitude; blue fill shades indicate DEGs upregulated in neonatal RTEs, and green fill shades indicate DEGs upregulated in adult RTEs,AII counted DEGs have a expression value greater than 20. (G) Bubble plot depicting the gene set enrichment results when comparing the per cluster age-related differential expression results to the virtual memory (VM), true naïve (TN), and ImmGen T cell gene signature (II, III, IV, VI, Vil, VIII, X) gene sets. Dot shading indicates the magnitude and direction of the normalized enrichment score (NES), while dot sizes indicate the adjusted p-value, with larger dots corresponding to smaller adjusted p-values. The “0-10” row displays the enrichment results calculated by comparing all neonatal RTE transcriptomes to all adult RTE transcriptomes.

To identify the transcriptional programs of individual clusters within each supercluster, we performed gene set enrichment analysis (GSEA) ^38^, querying genes distinctly up or downregulated within each cluster against ImmGen, KEGG, and ImmPort curated pathways ^39-41^. Within the effector-like supercluster (1 and 8), cluster 8 transcriptomes possessed particularly high enrichment for gene sets associated with an activated state, including “lymphocyte activation”, “initial cytokine or effector response (I)”, and “glycine, serine, and threonine metabolism” (Fig. S2A-C). Within the true naïve supercluster (0, 3, 5, 6 and 9), cluster 3 transcriptomes were distinctly enriched for the “memory precursor (VII)” gene signature; notably, enrichments were driven by the upregulation of the memory-associated genes *Il7r, Tcf7*, and *Bcl2* ^41^ (Fig. S2B). Interestingly, despite their overall true naïve transcriptional phenotype, clusters 5, 6, and 9 all expressed genes signifying a response to external stimuli. Cluster 5 transcriptomes were enriched for the “interferon alpha/beta signaling” gene signature, corresponding to the recently defined ISAG^hi^ (Interferon Signaling-Associated Genes) subset ^42,43^ (Fig S2C). Cluster 6 transcriptomes were uniquely enriched for NF-kB dependent pathways, such as “toll-like receptor cascades” and “T cell activation”, possessing an upregulation of NF-kB subunits (*N>b2, Relb*) and targets (*Bcl3, Icam1*) (Fig. S2C). Cluster 9 transcriptomes were enriched for “steroid biosynthesis”, potentially indicating an effector-primed metabolism, given the upregulation of cholesterol production following T cell activation^44^ (Fig. S2A). Within the virtual memory supercluster (2, 4, 7, and 10), cluster 7 uniquely expressed high levels of the Treg-associated transcripts *Izumo1r, Ikzf2*, and *Ctla4* (Fig. 2B). Finally, while clusters 4 and 10 shared enrichment for the “short term effector and memory (VI)” gene signature, cluster 10 transcriptomes were potentially analogous to a recently described human subset of NK-like CD8+ γδ T cells, a possibility suggested by the distinct enrichment of multiple gene sets associated with NK cell function and high γδ TCR gene set module scores ^45,46^ (Fig. S2B-C).

Having characterized the phenotypic composition of the RTE pool, we next asked how the neonatal and adult RTE pools differed, first hypothesizing that the RTE pools could differ because of age-related differences in the proportion of cells from each of the defined clusters. To address this possibility, within each cluster, we compared the proportion of cells derived from neonates to the proportion of cells derived from adults. We found that neonatal RTEs contributed more to the virtual memory clusters 2 and 4, whereas adult RTEs contributed more to the true naïve clusters 5 and 9 (Fig. 2D-E). Therefore, neonatal and adult RTE pools may be biased towards different infection responses due to differences in their phenotypic composition in the steady state.

In addition to differences in the cluster composition of the RTE pool, age-related differences in gene expression within each cluster could also impact how the neonatal and adult RTE pools respond to infection. Indeed, neonatal and adult RTE pools also possessed core transcriptomic differences; by performing differential expression analysis to compare neonatal and adult transcriptomes on a per-cluster basis, we identified 897 differentially expressed genes (Fig. 2F). To determine whether the age-related differences in gene expression were either consistent across the defined clusters or cluster-specific, we first employed a k-means clustering approach, which revealed 6 clusters of genes differentially expressed by age. Within each of the identified k-means clusters, the differentially expressed genes were either upregulated across nearly all phenotypic clusters in neonatal RTEs (K1, K2, K6) or upregulated across nearly all phenotypic clusters in adult RTEs (K3, K4, K5) (Fig. S2D). To gain insight into the functional significance of the core transcriptomic differences between neonatal and adult RTEs, GSEA was performed using the per-cluster, age-related differential expression results. The analysis revealed positive enrichments (indicating upregulation in neonatal RTEs) for the “virtual memory”, “preparation for cell division (II)”, “late effector or memory (X)”, “short-term effector and memory (VI)”, and “cell cycle and division (III)” gene signatures. Conversely, the “true naïve”, “naïve and late memory (IV)”, “memory precursor (VII)”, and “naïve or late effector or memory (VIII)” gene signatures were negatively enriched, indicating upregulation of their constituent genes in adult RTEs (Fig. 2G). The consistency of the enrichment results across the phenotypic clusters demonstrates that the neonatal RTE pool not only possesses more cells with a virtual memory phenotype, but that all neonatal RTEs, regardless of their phenotype, possess a more effector-biased transcriptome than adult RTEs.

### TCR usage in neonatal and adult CD8+ RTEs

We next sought to determine the key factors that contribute to the altered gene expression profiles between neonatal and adult RTEs. For example, why do neonatal CD8+ RTEs exhibit a more VM phenotype? One possibility is that the altered developmental trajectory among neonatal RTEs corresponds to their usage of different T cell receptors (TCRs). To examine this possibility, we analyzed the TCR data in our paired scTCR/RNA-seq datasets for neonatal and adult RTEs. We first asked whether TCR composition was skewed in CD8+ RTEs produced in early life. When we compared V gene usage, we observed a similar pattern of V genes expressed in both the TCRβ and TCRα chains of RTEs produced in neonatal and adult mice (Fig. S3A-B). We did not observe a significant difference in J gene usage in neonatal and adult CD8+ RTEs, another similarity between the two groups (Fig. S3C-D). Thus, despite their distinct phenotypes, the usage of V and J genes were largely comparable in both neonatal and adult CD8+ RTEs.

For a more in-depth comparison, we next focused on particular features in the CDR3 regions of the TCRβ and TCRα chains. Previous work has demonstrated that neonatal T cells have more germline-encoded TCRs (with no N-nucleotide additions) compared to adults, as expression of TdT (the enzyme responsible for insertion of N-nucleotides in the junctional regions) is not expressed in the thymus of mice until 4-8 days of age ^47,48^. Consistent with these studies, we found that the timestamped CD8+ T cells in neonatal mice showed a significantly higher proportion of cells with no N-additions in the TCRβ chain compared to their counterparts in adult mice (Fig. S3E). The lack of junctional diversity in neonatal CD8+ RTEs also translated into shorter CDR3β lengths. In contrast, we did not observe significant age-related differences in the numbers of N-additions in the TCRα chain (Fig. S3F), which is consistent with earlier work ^49^. We also found a smaller number of unique clonotypes in neonatal CD8+ RTEs (Fig. SG), suggesting that the altered phenotype of CD8+ T cells made in early life corresponds to a less diverse pool of T cells, which is biased towards shorter and more germline-encoded TCRs.

We next sought to determine whether the bias towards VM cells is linked to the usage of more germline-encoded TCRs. To address this question, we compared the proportion of cells with germline encoded TCRs across each gene expression cluster and found that the germline-encoded RTEs were slightly more prevalent in clusters 4 (neonatal and adult) and 10 (neonate only), both of which possess a VM phenotype (Fig 3A). Further, to ask whether neonatal RTEs that express TCRs with zero N-additions possess a heightened effector-like phenotype, we performed differential expression analysis comparing VM RTEs with zero N-additions to neonatal VM RTEs with >2 N-additions (Fig. 3B-C) and found that neonatal VM cells with germline-encoded TCRs upregulated genes (*Cxcr3, Ly6c2, Il2rb*) that are associated with the VM phenotype (Fig. 3B). GSEA revealed that the “late effector or memory (X)”, “virtual memory”, “cytokine - cytokine receptor interaction”, and “memory precursor (VII)” gene signatures were positively enriched, indicating a coordinated upregulation of T cell response-associated genes (*Ccl5, Gpr183, Tbx21*) within neonatal cells possessing germline-encoded TCRs (Fig. 3D-E). Although these differences may be driven by a small effect size, we did not observe differentially expressed genes or strong gene set enrichment results when comparing the gene expression profiles of adult RTEs with zero N-additions versus those with >2 N-additions (Fig. 3B; S3H), indicating germline-encoded TCRs only exhibit a distinct transcriptional signature in early life. Collectively, these data suggest that neonatal CD8+ RTEs with the most effector-like gene expression profiles tend to express germline-encoded TCRs.

**Figure 3.**
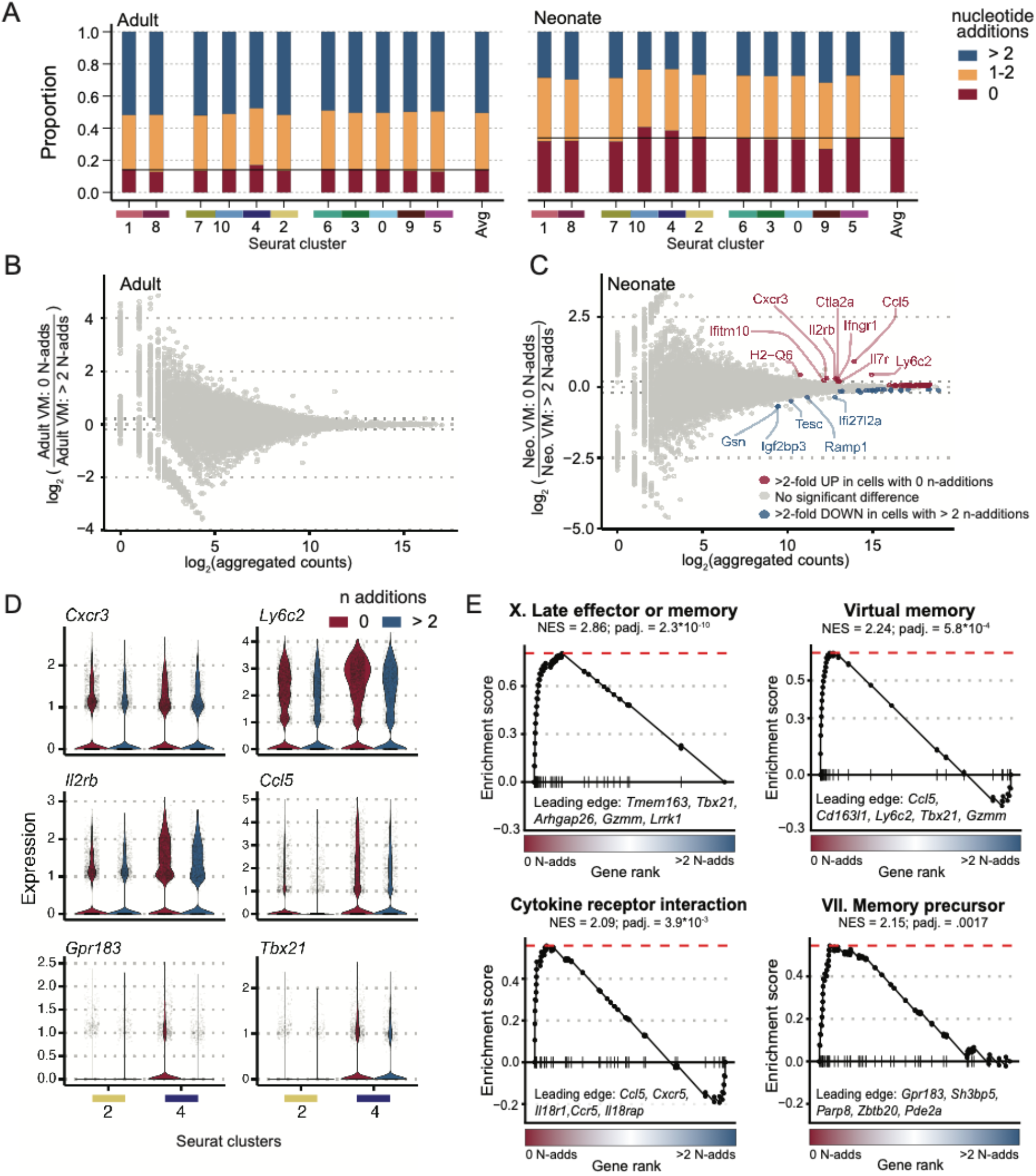
TCR complexity is associated with the effector-biased transcriptome of neonatal RTEs. (A) Proportion of RTEs with different numbers of TCRB N-nucleotide additions. Proportions are calculated after cells are grouped by gene expression clusters and age. Fill color indicates the number of N-additions binned into 0, 1-2, and >2 groups. (B- C) MA plots visualizing the relationship between gene expression (x-axis) and log_2_(fold-change) when comparing either adult (B) or neonatal (C) RTEs that possess a virtual memory transcriptome (gene expression clusters 2 and 4) and TCRBs with 0 N-additions to RTEs that possess a virtual memory transcriptome and TCRBs with > 2 N-additions. Each point represents a gene, and point color represents the significance and direction of the differential expression result. All red and blue points have an adjusted p-value of < 0.05, while labeled points have a log_2_(fold-change) with an absolute value of > 0.20. (D) Violin plots showing the distribution of single-cell gene expression (y-axis) for Cxcr3, Ly6c2, Il2rb, Ccl5, Gpr183, and Tbx21. Single cell expression values are grouped along the x-axis by gene expression cluster (columns labeled 2 and 4) and number of N-additions (red = “0” and blue = “> 2”). Gpr183 and Tbx21 are not statistically significantly differentially expressed, but they appear within the leading edge of enriched pathways in (E). (E) GSEA running enrichment score plots for four selected pathways significantly upregulated in neonatal virtual memory phenotype RTEs with 0 N-addition TCRBs compared with > 2 N-addition TCRBs [gene rank expressed as mean log2(Neo. VM: 0 N-adds/ Neo. VM: ≥ 2 N-adds)]. NES, normalized enrichment score.

### Functions of neonatal and adult RTEs

An important question is whether neonatal and adult RTEs exhibit distinct functional properties. We previously showed that neonatal CD8+ T cells undergo more cell divisions than adult CD8+ T cells after *in vitro* TCR stimulation and preferentially give rise to short-lived effectors (KLRG1+ CD127) after *in vivo* infection ^9-11,50^. However, these experiments were performed with bulk CD8+ T cells from neonatal and adult animals, raising the possibility that the increased percentage of RTEs in the neonatal pool may be underlying these age-related differences. To examine this possibility, we again used our timestamp mice to generate neonatal and adult CD8+ RTEs. This time, we coated the neonatal and adult RTEs with proliferation dye (CellTrace Violet, CTV) and compared their ability to proliferate after *in vitro* stimulation with plate-bound anti-CD3/anti-CD28 (Fig. 4A). Two days after stimulation, we examined the dilution of CTV, a proliferation metric, and found that neonatal RTEs had undergone more divisions than adult RTEs (Fig. 4B-C). Thus, even after controlling for age-related differences in post-thymic maturation, neonatal CD8+ T cells were still more proliferative than adult CD8+ T cells.

**Figure 4.**
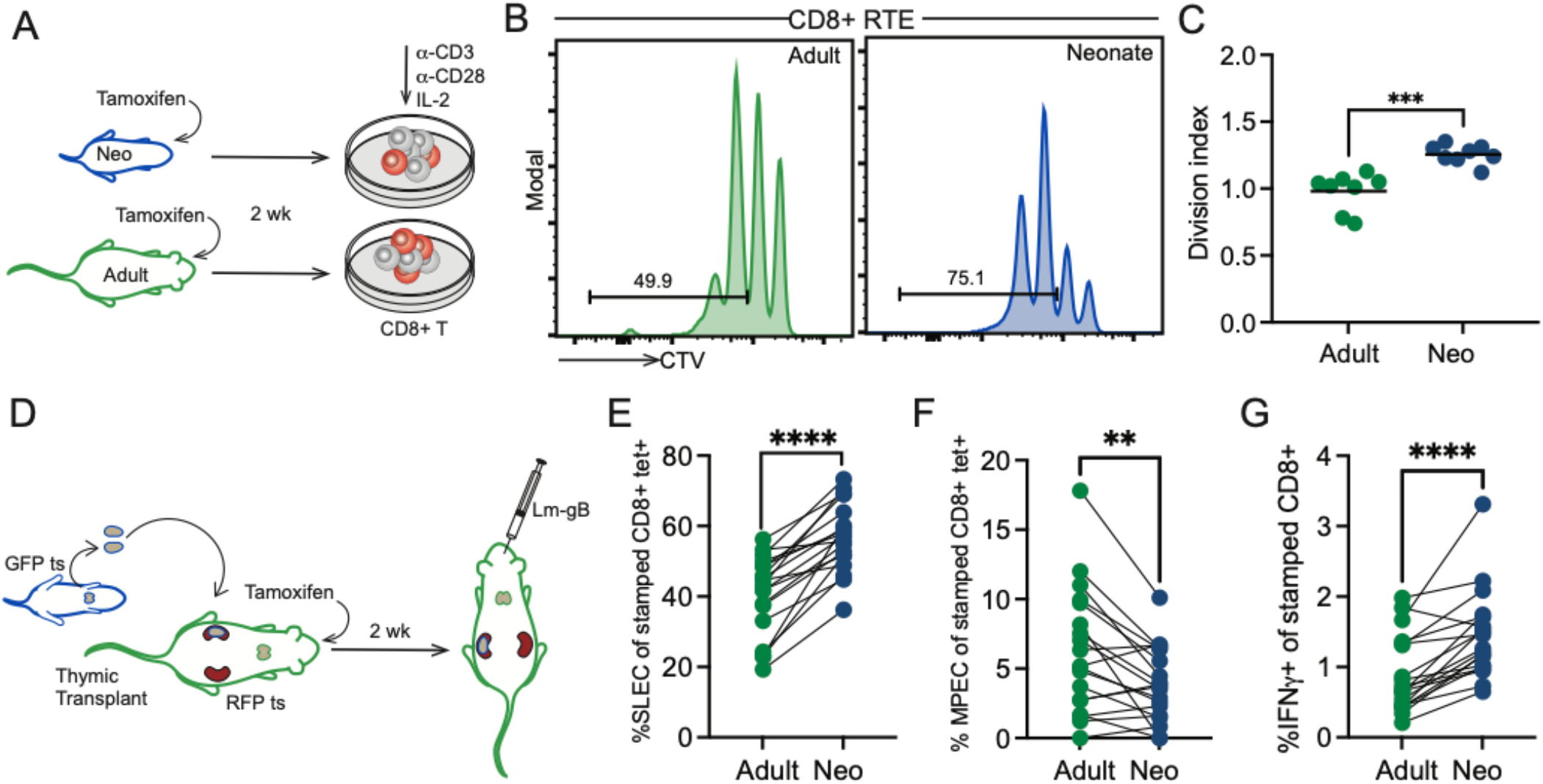
Neonatal RTEs exhibit enhanced immune functionality. (A) Schematic of in vitro stimulation. CD8+ T cells from neonatal and adult timestamp mice were coated with cell trace violet (CTV) and stimulated for 48 hrs via antibody- mediated crosslinking of CD3/CD28. (B) Representative histograms of CTV dilution. (C) Division index. Unpaired t-test was performed for statistical analysis. N=8 from 2 independent experiments. (D) Schematic of in vivo infection experiment. Thymic lobes from newborn GPF timestamp mice were grafted under the kidney capsule of adult red timestamp mice. Tamoxifen was administered to thymic graft recipients, and after 2 weeks, recipients were infected with 5x10^3^ CFU L. monocytogenes-gB. Spleens were collected at 5 dpi. (E) Percentage of timestamped tetramer positive cells with a short-lived effector phenotype (KLRG1^lo^ ’CD127^hi^). (F) Percentage of timestamped tetramer positive cells with a memory precursor effector phenotype (KLRG1^lo^CD127^hi^). (G) Percentage of timestamped cells producing IFN γ. N=19. For statistical analysis, paired t-tests were performed. **, P<0.01; ^***^, P<0.001; ^****^, P<0.0001./

To control for age-related differences in the peripheral environment, we repeated our thymic transplant experiments. For these studies, we administered tamoxifen after thymic transplantation to generate waves of neonatal (ZsGreen) and adult (TdTomato) CD8+ RTEs in the same host (Fig. S4A). Importantly, neonatal CD8+ RTEs still underwent more proliferation than adult RTEs after *in vitro* TCR stimulation (Fig. S4B-C). We also infected the dual timestamp thymic transplant mice with a recombinant strain of *Listeria monocytogenes* that expresses the gB peptide from HSV-1 (denoted LM-gB; Fig. 4D) and compared the ability of neonatal and adult RTEs to differentiate into effector cells. Strikingly, a higher proportion of neonatal RTEs displayed a short-lived effector phenotype (SLEC; Fig. 4E) and fewer memory precursor effector cells (MPEC; Fig. 4F). The neonatal RTEs also produced more effector molecules than adult RTEs, such as IFNγ (Fig. 4G), granzyme A, granzyme B, and TNFα (Fig. S4D-F), after peptide stimulation. In contrast, the adult RTEs generated fewer SLECs and more MPECs. Collectively, these data suggest that post-thymic maturation contributes very little to differences between neonatal and adult CD8+ T cells; instead, such differences are largely cell intrinsic.

## Discussion

In this report, we leveraged a new tool to identify RTEs in neonatal and adult animals, allowing us to directly compare their phenotype and functions. We found that neonatal CD8+ RTEs are distinguished from adult CD8+ RTEs by their expression of distinct surface markers, gene expression profiles, and TCR features. The neonatal CD8+ RTEs also respond more rapidly to stimulation than adult CD8+ RTEs and give rise to more terminally differentiated effector CD8+ T cells after infection. These data indicate that RTEs are made differently in early life and that post-thymic maturation is not a major driver of cell-intrinsic differences between neonatal and adult CD8+ T cells.

In addition to ‘timestamp’ mice, we used paired TCR/RNA-sequencing to construct single cell atlases of neonatal and adult RTEs. While many of the developmental-related differences in the RTE pool may be attributed to their derivation from distinct hematopoietic progenitors ^14,51^, there is still a surprising amount of heterogeneity within each RTE population for reasons that are unknown. Our data suggest that the TCR is a factor that contributes to phenotypic variation in the neonatal CD8+ RTE pool. Consistent with previous studies, we found that neonatal CD8+ T cells exhibit shorter and more germline-encoded TCRs ^52,53^. However, by simultaneously examining gene expression and TCR sequences in individual cells, we discovered that germline-encoded TCRs exhibit a bias towards becoming virtual memory cells in early life.

Although it is not clear why neonatal RTEs with germline-encoded TCRs have a propensity to become virtual memory cells, several possibilities are worth mentioning. First, germline-encoded TCRs have been shown to be more cross-reactive, or ‘peptide-promiscuous’ ^54^, which may enable them to receive stronger signaling in response to self-peptide:MHC complexes in the periphery. Second, TCRs with more N-additions tend to be longer, and longer TCRs may impair MHC binding and TCR signaling in the thymus ^55^. Third, the germline-encoded TCR may correspond to an early wave of hematopoietic progenitors that are programmed differently. Indeed, recent studies have indicated that T cells can be produced by hematopoietic progenitors that arise from endothelial cells in the yolk sac and embryo proper (prior to the emergence of HSCs) ^56,57^. However, the TCR repertoire of such HSC-independent CD8+ T cells has yet to be explored.

Previous studies have uncovered distinct transcriptional states in the naïve CD4 pool, including some clusters that are distinguished by their expression of TCR pathway genes, memory-like genes, or interferon response genes ^58,59^. Interestingly, we observed similar clusters in the CD8+ RTE pool, which are preferentially populated by cells made at different stages of life. The CD8+ T cells produced in neonatal mice have a propensity to express more memory-like genes, whereas those made in adult mice give rise to the subsets that are enriched for interferon responsiveness and cholesterol biosynthesis. By using timestamp mice, we were able to compare CD8+ T cells that were definitively ‘neonatal’ or ‘adult’ and demonstrate how the time of production contributes to transcriptional heterogeneity in the naïve T cell pool.

In the future, it will be important to determine which cell subsets and TCRs are preferentially preserved in the adult T cell pool. Indeed, we previously demonstrated that some (but not all) neonatal CD8+ T cells persist into adulthood, and the ones that persist are the first CD8+ T cells to respond to infection. However, the transcriptional and clonotypic features of these long-term survivors have remained a mystery. Similarly, we do not yet know which transcriptional and TCR features are present on the neonatal cells that preferentially give rise to short-lived effectors in adult animals. While these questions remain to be answered, this study provides new insight into how CD8+ T cells are made differently at various stages of life.

### Materials and Methods Mice

ZsGreen (#007906), TdTomato (#007909), and TCRδCre-ERT2 (#031679) mice were obtained from Jackson Laboratories. All experiments utilized timed matings to ensure mice were of similar ages. Mice were maintained under specific pathogen-free conditions at the College of Veterinary Medicine. The facilities are accredited by the American Association of Accreditation of Laboratory Animal Care. All protocols regarding animal use were reviewed and approved by the Institutional Animal Care and Use Committee at Cornell University.

### Timestamping method

Timestamp mice were generated by crossing CRδCre-ERT2 mice with ZsGreen or TdTomato reporter mice. To activate fluorescent reporter expression in neonatal T cells, 2.5 mgs of tamoxifen were administered to dams by oral gavage three times in 12-hour intervals over a 24-hour period and 0-1d old pups received tamoxifen through lactation. To activate fluorescent reporter expression in adult-derived T cells, 1 mg was administered to 4 wk old mice by oral gavage in 24-hour intervals for two days.

### Thymic transplants

Thymic transplants were performed as previously described^14^. Briefly, thymus lobes were isolated from 0-1d old timestamp mice. The thymus lobes were separated into individual lobes and were placed under the kidney capsule of a 6-week-old mouse. To mark thymocytes from both donor and recipient, 5 mg of tamoxifen was administered from 0-3d post-transplantation. Recipients were sacrificed at indicated times and spleens were isolated to assess phenotype and behavior of stamped cells through flow cytometry.

### Listeria Infections

Mice were infected with 5×10^3^ CFU of wild-type (WT) Listeria monocytogenes expressing the gB-8p peptide (Lm-gB) as previously described ^12^.

### Flow cytometry

For flow cytometry, cells were stained with antibodies purchased from ThermoFisher/Life technologies, BioLegend, or BD Biosciences. The following clones were used: CD4 (GK1.5), CD8a (53-6.7), Qa2 (1-1-2), CD103 (2E7), CD127 (A7R34), CD11a (M17/4), KLRG1 (2F1), IFNγ (XMG1.2), granzyme A (GzA-3G8.5), granzyme B (GB11), Ki76 (SolA15) and Eomes (Dan11mag). Viability staining was done using Fixable Viability dye e780 (ThermoFisher/Life technologies). For tetramer staining, biotinylated monomer was obtained from the NIH Tetramer core facility and tetramerize with APC-Streptavidin (ThermoFisher/Life technologies). For surface staining only, IC fix buffer set (ThermoFisher/Life technologies) and for experiments needing intracellular staining, BD Cytofix/Cytoperm fixation kit was used, both according to manufacturer’s instructions. Flow cytofluorimetric data were acquired using FACSDiva software from a BD FACSymphony A3 or A5 SE, both equipped with five lasers (BD Biosciences). Analysis was performed with FlowJo (BD Biosciences).

### *In vitro* stimulation

CD8+ T cells were isolated from the spleen using positive magnetic selection with CD8 microbeads (Miltenyi). Following magnetic bead purification, cells were labeled with Cell Trace Violet (ThermoFisher/Life technologies) according to manufacturer’s recommendation. Cells were stimulated with plate-bound anti-CD3 (2.5 ug/ml, clone 2C11) then cultured with complete RPMI supplemented with 100 U/ml human IL2 and 4 ug CD28/ml (clone 37.51). Cells were harvested at 48 hrs, stained for surface markers and then analyzed by flow cytometry.

### Flow cytometry statistical analysis

Statistical analysis was performed using Prism (Graphpad). Error bars represent SEM. Statistical significance was determined by paired or unpaired student’s t-test as indicated in figure legends. Significance for individual figures is denoted in the legend.

### Single-cell RNA-seq

#### scRNA-seq library preparation

Two weeks post timestamp, live, splenic ZsGreen+Va2+CD8+ cells were sorted into 0.04% BSA in PBS. Cells were loaded in the Chromium instrument (10X Genomics) for the formation of gel bead-in-emulsions (GEMs) targeting 5-10k cells per sample. Single-cell RNA-seq and TCR libraries were prepared using Chromium Single Cell Universal V(D)J Reagent Kits (v1.1 chemistry; 10X Genomics) by the BRC Genomics Facility following the manufacturer’s protocol. Libraries were sequenced on a NovaSeq 6000 (Illumina) to an average depth of >150M reads (scRNA-seq) and >15M reads (scTCR).

#### scRNA-seq raw data processing and cell quality filtering

Raw FASTQ data was processed using ‘cell ranger multi’ (v6.0.0, 10X Genomics) with provided mouse reference genome refdata-gex-mm10-2020-A and TCR reference refdata-cellranger-vdj-GRCm38-alts-ensembl-5.0.0 for TCRs. The resulting count matrixes were filtered using Seurat v5.0.0 to remove cell barcodes with 1) fewer than 500 genes detected, 2) fewer than 1000 transcripts detected, 3) more than 7.5% mitochondrial counts, and 4) those not passing TCR sequence filtering. After filtering, we obtained high-quality transcriptomic profiles for 59,281 individual cells with an average of 7,410 cells per sample and 1,560 genes and 4,419 transcripts detected per cell.

#### scRNA-seq dataset preprocessing, identity-based filtering, and clustering

Following cell quality filtering, three rounds of data preprocessing were performed. Each preprocessing round included: count normalization, scaling, variance regression, dimensionality reduction, cluster identification, and UMAP generation. Within each preprocessing round, we specifically regressed out variation in mitochondrial read percentage, ribosomal read percentage, cell cycle associated gene expression, and TCR gene segment gene expression. Following an initial round of preprocessing, three clusters of cells were removed from the dataset, which corresponded to 1) library processing stress, 2) dividing cells, and 3) CD8+ DCs. After a second round of preprocessing was performed, a small number of cells associated with cell division were again identified; these cells were removed from the dataset. Upon a final round of preprocessing, we identified 11 total clusters (resolution of 0.7) that served as a basis for the downstream analysis.

#### scRNA-seq cluster annotation

Seurat’s FindMarkers function was used to compare each individual cluster of cells to the rest of the dataset, resulting in the identification of the 5 genes that were most upregulated per cluster. Bubble plot visualization of gene expression was generated with the Clustered_DotPlot function of scCustomize v2.1.2.9029. The lower and upper bounds of the bubble plot expression color scale were defined as the 10^th^ percentile expression value and 90^th^ percentile expression value considering the expression of all input genes across all clusters. Gene set module scores were calculated using the AddModuleScore_UCell function of UCell v2.6.2. The true naïve and virtual memory gene sets were derived from ^14^, ImmGen T cell gene signature gene sets are derived from^41^. GSEA was performed using fgsea v1.28.0 ^60^ comparing the gene sets against genes ranked by their fold-change as reported by Seurat’s find markers function. GSEA bubble plots were generated using ggplot2 v3.5.1.

#### scRNA-seq differential expression analysis

For differential expression analysis, we generated pseudo-bulk expression profiles for each cluster by aggregating the raw counts across single cells with the same sample and cluster identity using the aggregate.Matrix command of Matrix.utils v0.9.7. The resulting pseudo-bulk expression profiles were then used as input into DESeq2’s differential expression pipeline ^61^. Genes that possessed ≥ 3 counts across ≥ 3 pseudo-bulk samples per cluster were retained for differential expression testing. Age-related differential expression testing was performed within each cluster, and genes with an adjusted p-value less than 0.05 were considered to be differentially expressed. GSEA was performed per cluster by comparing gene sets against genes ranked by their DESeq2 reported fold-changes (neonate vs. adult). Finally, differentially expressed genes were clustered using the K-means clustering algorithm implemented by the Stats package v4.3.2 considering their fold-change values across each cluster. Six K-means clusters were plotted given the results of plotting the within-cluster sum of squares using the function fvis_nbclust within the factoextra package v1.0.7. Fold-change boxplots were generated using ggplot2.

#### Integrative scTCR-seq and scRNA-seq analysis

To integrate TCR information with single cell expression profiles, each cell was assigned an annotation of 0, 1-2, or ≥ 2 TCR n-additions. The average proportion of cells with 0, 1-2, and ≥2 N-addition annotations were calculated separately for adult and neonatal cells after first grouping cells by age of origin. Cluster-specific proportions of cells with 0, 1-2, and ≥2 N-addition annotations were calculated after cells were grouped by their age of origin and gene expression cluster. Differential expression testing was performed between cells with 0 and ≥ 2 N-additions after grouping cells by 1) age of origin, 2) age of origin and virtual memory phenotype (gene expression clusters two and four), and 3) age of origin and true naïve phenotype (gene expression clusters zero and three). Gene expression values were calculated for MA plot visualization by aggregating the normalized counts across each cell per gene, for all detected genes. Fold-change values reported were calculated via Seurat’s FindMarkers function. Genes were denoted as significantly differentially expressed if their associated adjusted p-values were less than 0.05. Violin plots were generated using normalized count data as input into Seurat’s VlnPlot function. GSEA was performed using fgsea v1.28.0 comparing the gene sets against genes ranked by their fold-change as reported by Seurat’s find markers function. GSEA bubble plots were generated using ggplot2 v3.5.1, and running enrichment score plots were generated with fgsea’s plotEnrichment function.

### TCR repertoire analysis

The full-length consensus TCR sequences assembled by Cell Ranger were aligned against the TCR genes using IMGT/HighV-QUEST^62^ to determine the best-matched V, (D), and J genes and the minimum number of nucleotide additions required to generate each TCR sequence. TCR sequences were omitted from subsequent analysis on the basis of non-productive rearrangements, unidentified V or J gene usage, and more than one productive alpha or beta chain. TCR diversity was assessed as the number of unique TCR clonotypes. The diversity analysis was restricted to cells for which there was a TRA-TRB sequence pair and cells were pooled across the 4 mice in each of the neonatal and adult groups. To account for differences in sample size, 12000 cells were randomly drawn from each pooled population and the number of unique TRA, TRB, and TRA-TRB clonotypes were estimated for each of 1000 random draws.

## Acknowledgments

We thank the Flow Cytometry Facility (RRID:SCR_021740) for expert sorting assistance and the BRC Genomics Facility (RRID:SCR_021727) for generation and sequencing of libraries. This work was supported by National Institute of Health awards R01HD107798, R01AI05265, R01AI110613 (to B.D.R, from the National Institute of Allergy and Infectious Disease), R21AI138025 (to N.L.S. from National Institute of Allergy and Infectious Disease) and F31AI157236 (to C.T., from National Institute of Allergy and Infectious Disease), and Australian National Health and Medical Research Council Career Development Fellowship 1067590 (VV).

