## Supplemental figures for "Single-cell multiomics reveals the distinct properties of neonatal and adult recent thymic emigrants"

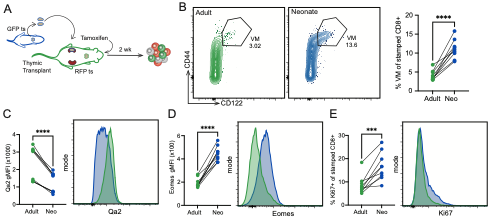

**Figure S1. Cell intrinsic differences between neonatal and adult RTEs.** (A) Schematic of experimental set up. Briefly, thymic lobes from newborn GFP timestamp mice were transplanted into 6wk old RFP timestamp mice. Recipient mice were given tamoxifen to label RTEs and mice were collected for cells at 2 weeks post marking. (B) Representative contour plots displaying virtual memory (VM, CD44^hi^ CD122^hi^) CD8+ populations (left) and statistical analysis of the VM population within the marked T cell population (right). (C) Statistical analysis and representative histograms of RTE marker Qa2. (D) Statistical analysis and representative histograms of Eomes expression. (E) Statistical analysis and representative histograms of Ki67 staining. N=8 from 2 independent experiments. For statistical analysis, Paired t-tests were performed. ***, P<0.001; ****, P<0.0001.

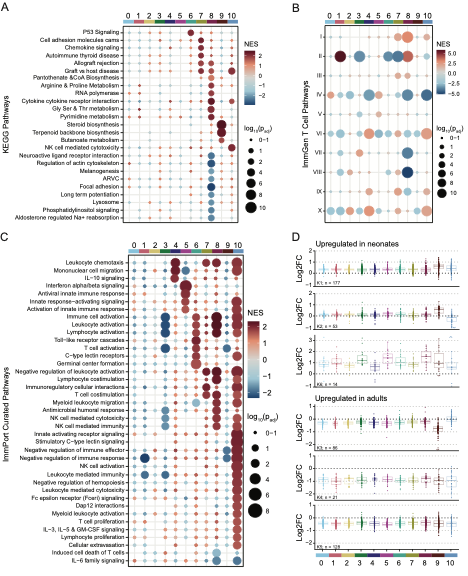

**Figure S2. Gene expression variation between neonatal and adult RTEs. (**A-C) Bubble plots depicting Normalized Enrichment Scores (NES) for all significantly enriched KEGG (A), ImmGen T cell (B), and Immport curated GO:BP and Reactome (C) pathways when ranking genes based on the fold-change derived from pairwise comparisons between each individual cluster vs all other clusters. Dot size indicates the adjusted p-value, where larger dot size indicates smaller values. (D) Boxplots depicting the log_2_(fold-change) of differentially expressed genes (DEGs) between neonatal and adult RTEs. Each panel corresponds to a distinct cluster of DEGs determined by the K-means clustering of DEGs by their log2(fold-change) values across the 11 phenotypic gene expression clusters, where n = # DEGs in each K-means cluster. Within each panel, the log2(fold-change) value for each DEG is plotted for all 11 clusters. The consistency across the 11 phenotypic clusters in each panel indicates that the genes are consistently differentially expressed between neonatal and adult RTEs irrespective of the phenotypic cluster identity.

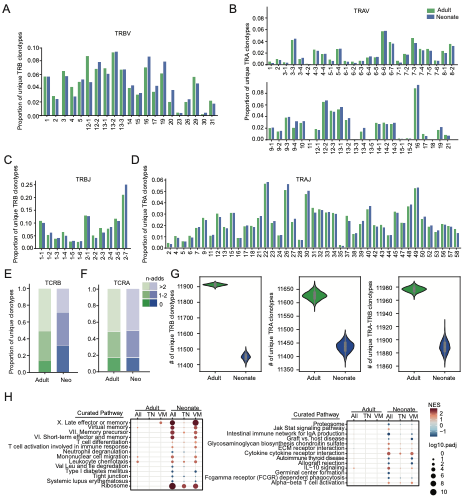

**Figure S3.** **TCR repertoire analysis of adult and neonatal RTEs.** (A-B) Individual V gene usage by adult and neonatal RTEs: (A) TRBV and (B) TRAV. (C-D) Individual J gene usage by adult and neonatal RTEs: (C) TRBJ and (D) TRAJ. V and J gene numbering corresponds to IMGT gene nomenclature and multiple best-matched V and J genes, with proportion <= 0.001, have been omitted. (E-F) Estimated minimal number of N-nucleotide additions for TCRB (E) and TCRA (F). (G) Distribution of the number of unique clonotypes by TCRB (left), TCRA (middle) or TCRA-TCRB pairs (right) estimated for 1000 random draws of a standardized sample size of 12000 cells per neonate and adult group. (H) Bubble plot depicting the gene set enrichment results when comparing differential expression results to the virtual memory (VM), true naïve (TN), and KEGG, ImmGen T cell, and ImmPort curated gene sets (rows). Columns indicate the group within which differential expression testing was performed between cells with 0 N-additions and >2 N-additions for TCRB. All indicates that all cells within a given age of origin were tested; TN indicates that cells with a true naïve phenotype (clusters 0 and 3) within a given age of origin were tested; VM indicates that cells with a virtual memory phenotype (clusters 2 and 4) were tested. Dot shading indicates the magnitude and direction of the normalized enrichment score (NES), while dot sizes indicate the adjusted p-value where larger dots correspond to smaller adjusted p-values.

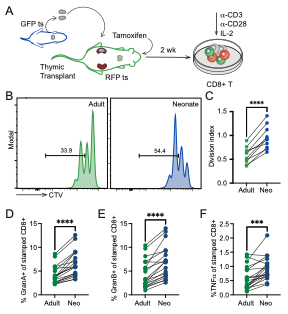

**Figure S4. Neonatal RTEs are more proliferative and cytotoxic.** (A) Schematic of *in vitro* stimulation. CD8+ T cells from thymic transplant timestamp mice were isolated 2wk post marking, coated with cell trace violet (CTV) and stimulated for 48 hr via antibody-mediated crosslinking of CD3/CD28. (B) Representative histograms of CTV dilution and (C) calculation of division index. For B-C, N=8 from 2 independent experiments. (D) percentage of timestamped cells with Granzyme A staining at 5dpi. (E) percentage of timestamped cells with Granzyme B staining at 5dpi. (F) percentage of timestamped cells producing TNFα at 5dpi. For D-F, N=19. For statistical analysis, paired t-tests were performed. ***, P<0.001; ****, P<0.0001.
